# SwitchClass: dissecting attenuated and escalated molecular features via a label-switch classification framework

**DOI:** 10.1101/2025.10.29.685265

**Authors:** Di Xiao, Pengyi Yang

## Abstract

Biological systems exhibit complex molecular trajectories in response to perturbations, ranging from changes that revert or attenuate towards homeostasis to alterations that persist or escalate. Capturing these complex patterns is essential for understanding molecular resilience and maladaptive persistence. Here, we introduce SwitchClass, a label-switch classification framework that distinguishes *attenuated* and *escalated* molecular features across biological states. By training dual classifiers with inverted outcome labels, SwitchClass computes a differential feature importance score (*δ*) that quantifies directional change in high-dimensional data. Applied to colorectal cancer proteomics spanning healthy, pre-treatment, and post-treatment samples, Switch-Class reveals proteins that normalise after therapy and those remaining dysregulated, uncovering partial molecular recovery. In phosphoproteomics of dietary perturbation and reversal, it uncovers phosphorylation sites linked to incomplete restoration of insulin signalling. In single-cell transcriptomes from COVID-19 patients with varying severities, it identifies cell-type-specific transcripts that mark either the resolution or persistence of inflammatory activity. Together, these analyses establish SwitchClass as a generalisable and interpretable framework for mapping directional molecular changes underlying adaptation, divergence, and disease severity across biological systems. SwitchClass is freely available from https://github.com/PYangLab/SwitchClass.

## Introduction

Cellular signalling and regulatory networks often respond to perturbations through molecular trajectories that can restore equilibrium or reinforce altered functional states (Kitano, 2004). At the molecular level, such perturbations remodel signalling and transcriptional networks in ways that determine whether cells remain close to a baseline configuration or diverge toward altered states. In cancers, large-scale proteomic analyses have revealed extensive molecular reprogramming in response to therapeutic intervention (Y. Li et al., 2024). Such changes capture divergent cellular trajectories, where subsets of molecular features normalise following treatment while others remain dysregulated, reflecting incomplete molecular recovery and treatment resistance. In metabolic systems, phosphoproteomic analyses have revealed extensive alterations in phospho-signalling under dietary perturbation, providing a foundation for examining how individual phosphorylation sites (phosphosite) align with normal or insulin-resistant signalling states (Fazakerley et al., 2023). In immune contexts, single-cell transcriptomic studies of COVID-19 have revealed marked heterogeneity in immune-cell activation across healthy, mild, and severe cases, highlighting shifts in interferon and inflammatory responses associated with disease severity (Schulte-Schrepping et al., 2020). These observations illustrate that biological perturbations rarely produce uniform molecular shifts. Instead, intermediate states frequently emerge, whether representing partial recovery or milder perturbation, where subsets of molecular features exhibit complex patterns that may align with baseline or perturbed states, collectively shaping the molecular configuration of the system. Yet, despite the prevalence of such multi-state systems, analytical frameworks capable of systematically quantifying these directional relationships across high-dimensional molecular features have remained limited.

Most existing approaches were developed to detect pairwise differences between conditions rather than to quantify directional relationships across multiple states. In transcriptomic and proteomic studies, frameworks such as limma-voom (Ritchie et al., 2015) and edgeR (Robinson, McCarthy, & Smyth, 2010) enable rigorous testing of group-specific effects in bulk data, while methods such as NEBULA (He et al., 2021) and muscat (Crowell et al., 2020) extend such pairwise analyses to single-cell transcriptomics. Although powerful for identifying statistically significant differences, these approaches are inherently limited to comparing one state against another. They provide little insight into how an intermediate condition (such as a dietary reversal, a partial treatment response, or a mild disease state) relates directionally to the baseline and the altered states. Trajectory-based frameworks such as tradeSeq (Van den Berge et al., 2020) and Monocle (Qiu et al., 2017) identify genes that vary continuously along inferred trajectories, but they assume ordered temporal progression, which is often inappropriate for cross-sectional or multi-condition datasets. While classification-based models and feature-selection algorithms can quantify feature importance (Yang, Huang, & Liu, 2021), they typically do so within a single label configuration and cannot reveal how feature relevance changes when relationships among states are inverted. As a result, current differential analysis strategies and feature selection methods offer limited insight into the directional behaviour of molecular features across multiple states, leaving the organisation of intermediate states largely unexplored.

To address this gap, we developed SwitchClass, a label-switch classification framework for quantifying directional molecular relationships across multi-state systems. SwitchClass builds on the idea that, given two extremes (e.g., baseline and strongly perturbed) and an intermediate state, each molecular feature can align more closely with one end. Practically, the framework constructs two complementary binary labellings of the data: (i) a baseline-aligned labelling, in which intermediate samples are grouped with the baseline and classified against the strongly perturbed state, and (ii) a perturbed-aligned labelling, in which intermediate samples are grouped with the perturbed state and classified against the baseline (Fig. 1A). A classifier is trained under each labelling, and a differential feature-importance score (*δ*) is then computed as the difference in feature relevance between the two models. The sign of *δ* encodes direction (attenuated if *δ >* 0, escalated if *δ <* 0), while |*δ*| reflects the strength of alignment. Because *δ* is defined through label switching, the framework is model-agnostic and can be implemented with any learning algorithm or feature importance metric.

**Figure 1.**
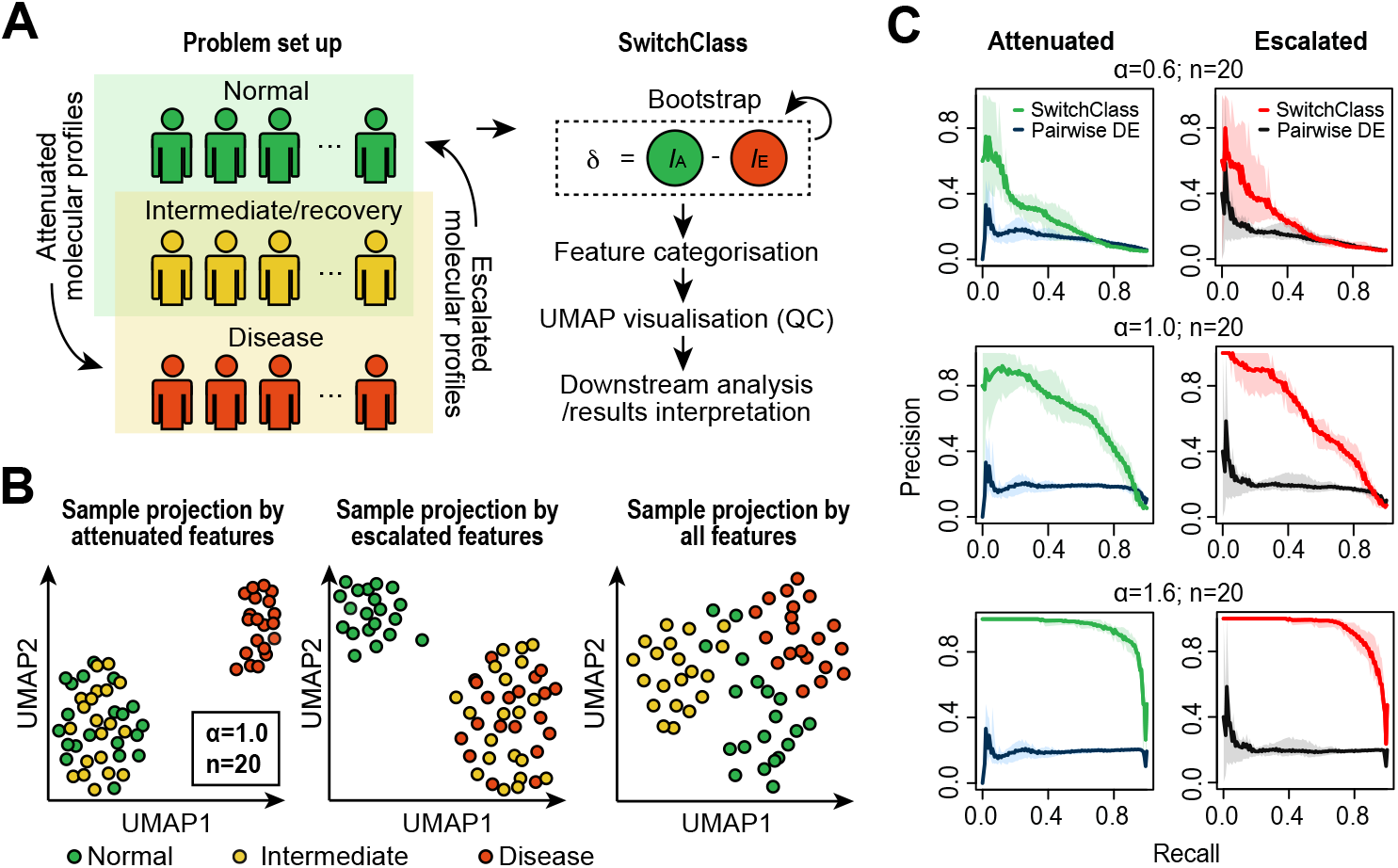
Conceptual overview and benchmarking of SwitchClass. (**A**) Schematic of the problem setup and the SwitchClass workflow. SwitchClass models molecular adaptation across three states (i.e., normal, intermediate/recovery, and disease) by quantifying directional importance scores (*δ* = *I*_A_ − *I*_E_) to classify features as attenuated or escalated. (**B**) UMAP projections of simulated samples using attenuated, escalated, and all features (*α* = 1.0; *n* = 20). (**C**) Precision-recall performance of SwitchClass in detecting attenuated and escalated features compared with pairwise differential expression (DE) across varying effect sizes (*α* = 0.6, 1.0, 1.6) and a sample size of *n* = 20 for each state. Shaded areas denote 95% confidence intervals derived from bootstrap resampling across simulation replicates.

Through simulations, we show that SwitchClass robustly detects directional signals across varying effect sizes and sample sizes. When applied to mass-spectrometry (MS)-based colorectal cancer proteomics spanning healthy controls and patient samples collected before and after treatment, SwitchClass reveals proteins whose abundances normalise following therapy and those that remain persistently dysregulated, implicating processes of partial molecular recovery and sustained tumor-associated activity. In mouse phosphoproteomics profiling dietary perturbation and reversal, the method identifies phosphorylation sites that normalise or remain elevated, uncovering signalling nodes linked to insulin resistance and its incomplete resolution. In single-cell transcriptomes spanning healthy individuals and mild and severe COVID-19 patients, SwitchClass identifies cell-type-specific transcripts in mild cases that either remain comparable to healthy individuals or are elevated to levels observed in severe disease. These patterns delineate distinct transcriptional states across multiple immune cell types that differentiate disease severity in response to COVID-19. Together, these analyses demonstrate the utility of SwitchClass and position it as a generalisable and interpretable framework for mapping directional molecular divergence across biological systems.

## Results

### Evaluating SwitchClass on simulated multi-state datasets

We first evaluated the performance and robustness of SwitchClass using simulated multi-state datasets where the true behaviour of each feature was known. Each dataset comprised three groups representing normal, intermediate, and disease states, corresponding conceptually to the baseline, intermediate, and perturbed conditions defined in the SwitchClass framework. A subset of 100 features was simulated to exhibit specific behaviour where for attenuated features, the intermediate group resembled the normal; for escalated features, it resembled the disease state. Additional features were generated to show either intermediate-specific deviations (200) or no change (700) (Supplementary Fig. 1A). The mean separation between normal and disease groups (effect size, *α*) and the number of samples per group (*n*) were systematically varied to test method performance under different signal and sample-size conditions (see Methods).

As expected, sample projections based on attenuated or escalated features reproduced the designed relationships among the three groups, in which UMAP embeddings derived from attenuated features grouped the normal and intermediate samples together while separating them from the disease group, whereas embeddings based on escalated features grouped the intermediate and disease samples and separated them from normal (Fig. 1B). In contrast, intermediate-specific features separated the intermediate group while null features produced no structured separation (Supplementary Fig. 1B).

Across simulation settings, SwitchClass recovered the expected directional behaviour of features, yielding *δ* scores that separate across the simulated categories (i.e., attenuated, escalated, intermediate-specific and null; Supplementary Fig. 1C). Specifically, attenuated features showed positive *δ* scores and escalated features showed negative *δ* scores, whereas intermediate-specific and null features were centred near zero. The magnitude of *δ* scaled with both the effect size (*α*) and the sample size (*n*), reflecting improved directional confidence as the contrast between normal and disease states strengthened. This behaviour demonstrates that the differential feature-importance score generated by SwitchClass faithfully encodes both the direction and strength of the alignment across multi-state systems.

To benchmark performance, we compared SwitchClass with a pairwise DE approach that tested normal against intermediate samples for escalated features and intermediate against disease samples for attenuated features, separately, on the above simulated datasets. Because this pairwise strategy evaluates only one boundary at a time and cannot integrate information across all three states, its ability to resolve directional relationships was limited, particularly when intermediate-specific effects were present. In contrast, SwitchClass achieved substantially higher precision-recall performance across all simulation settings as demonstrated by the higher precision-recall curves (Fig. 1C, Supplementary Fig. 1D). Notably, even under weak signal settings (*α* = 0.6), SwitchClass maintained relatively high precision, demonstrating that the label-switch design effectively leverages information across states to enhance directional inference. Together, these simulation experiments demonstrate that SwitchClass provides a reliable and interpretable measure of directional behaviour of features across multi-state systems, establishing a robust foundation for its application to empirical omic datasets.

### SwitchClass reveals attenuated and persistent proteomic signatures in colorectal cancer treatment

We applied SwitchClass to the plasma proteome dataset from a colorectal cancer cohort comprising healthy controls, pre-treatment, and post-treatment samples (Fig. 2A) (Y. Li et al., 2024). This configuration captures molecular trajectories from homeostasis through disease perturbation to partial therapeutic recovery. By computing differential feature-importance scores (*δ*) under dual label configurations, SwitchClass quantified for each protein whether post-treatment abundance aligned toward the healthy or disease state, classifying features as attenuated (*δ >* 0) or escalated (*δ <* 0) (Fig. 2A, Supplementary Fig. 2A). The resulting projections separated samples by health and treatment status, indicating a coherent directional structure within the plasma proteome (Fig. 2B).

**Figure 2.**
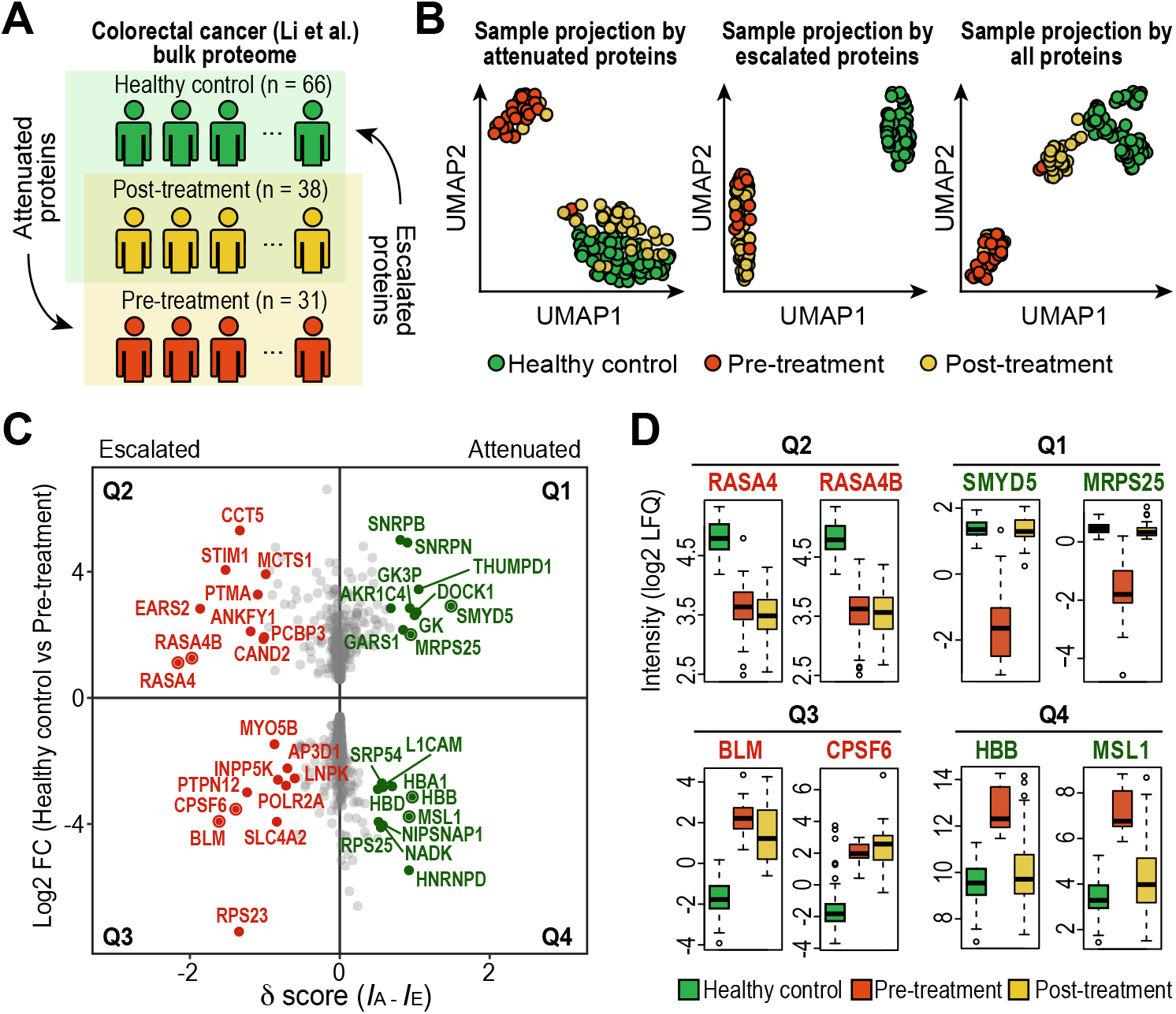
SwitchClass maps directional plasma proteomic changes across colorectal cancer progression and treatment. (**A**) Schematic overview of plasma sample collection from healthy controls, pre-treatment, and post-treatment colorectal cancer patients (Y. Li et al., 2024). (**B**) UMAP visualisation of sample projections using attenuated, escalated identified by SwitchClass, or all proteins. (**C**) Scatterplot of proteins based on log2 fold change (Healthy control vs Pre-treatment) and differential importance scores (*δ*) calculated by SwitchClass. Proteins with high |*δ*| in each of the four quadrants are highlighted. (**D**) Representative proteins from each quadrant illustrating attenuated (Q1/Q4) and escalated (Q2/Q3) proteins with distinct recovery or persistence profiles.

Attenuated proteins were enriched in factors involved in mitochondrial function, epigenetic regulation, and tissue homeostasis. Representative attenuated proteins across quadrants Q1 and Q4 include MRPS25 and SMYD5 (Q1), as well as HBB and MSL1 (Q4) (Fig. 2C, D). MRPS25, a mitochondrial ribosomal subunit required for OXPHOS protein synthesis, returned to healthy-like levels following treatment. This shift suggests a metabolic transition from glycolysis towards oxidative phosphorylation, a hallmark of recovered energy metabolism in colorectal cancer (Vander Heiden, Cantley, & Thompson, 2009). Similarly, SMYD5 functions as a tumor suppressor in colorectal cancer, in part through its role in chromatin regulation. Its depletion accelerates tumor growth and induces gene expression profiles resembling human colorectal cancer (Kidder et al., 2017). Restored SMYD5 expression after treatment indicates re-establishment of epigenetic regulation, particularly heterochromatin integrity. Elevated HBB levels in plasma are associated with heme-related oxidative stress, a common feature in cancer (Basak, Uddin, & Hancock, 2020). Their decline after treatment suggests reduced redox stress and a shift toward metabolic recovery. Finally, the recovery of MSL1, a component of the MOF histone acetylation complex, points to a reversal of chromatin hyperactivation and reinstatement of balanced histone modification activity (Luo et al., 2025). Collectively, these attenuated proteins reflect coordinated recovery of metabolic, epigenetic, and microenvironmental functions following therapy. Pathway enrichment analysis confirmed functional distinctions across quadrants (Supplementary Fig. 2B). Q1 proteins were enriched for chromatin regulation, histone methylation, and proteostasis control, while Q4 proteins were enriched for oxidative stress detoxification, chemical stress response, and amino acid metabolism. These signatures collectively point to systemic rebalancing of metabolic and epigenetic homeostasis as a feature of partial molecular remission.

Escalated proteins, by contrast, captured pathways and processes that remained pathologically active in post-treatment tumors. Proteins such as RASA4 and RASA4B (Q2) remained significantly down-regulated in both pre- and post-treatment tumors relative to healthy controls, indicating sustained loss of Ras GTPase activity and thus persistent RAS/MAPK pathway activation, thereby promoting malignant progression and drug resistance (X. Li et al., 2025). Enrichment analysis identified complement and coagulation cascades, heme-scavenging, and scavenger receptor activity among Q2 proteins, consistent with ongoing inflammation and disrupted metabolic clearance mechanisms (Supplementary Fig. 2B). Additional escalated responses were observed for BLM and CPSF6 (Q3) (Fig. 2C). BLM, involved in DNA repair, remained highly expressed after treatment, suggesting an ongoing DNA damage response or replicative stress in residual tumor cells. Clinically, elevated BLM levels are linked to poorly differentiated CRC subtypes and shorter relapse-free survival, highlighting its association with aggressive tumor phenotypes (Votino et al., 2017). Likewise, CPSF6, a core regulator of mRNA processing, remained persistently elevated, reflecting continued high transcriptional activity and cell proliferation demands despite treatment. Its sustained expression is linked to poor prognosis and treatment resistance in CRC, while knockdown restores drug sensitivity and limits tumor growth (C. Wang et al., 2025). Q3 proteins were enriched for receptor tyrosine kinase signaling and intracellular stress adaptation pathways, reflecting persistent survival signaling and resistance to molecular resolution.

Together, these results demonstrate that therapeutic intervention in colorectal cancer elicits a heterogeneous proteomic response. By separating proteins that normalise with treatment from those that remain persistently altered, SwitchClass captures the directional asymmetry of the treatment response and provides an interpretable framework for resolving how recovered and resistant molecular features coexist in treated tumours. This approach delineates a proteomic basis for partial remission, identifying candidate pathways (e.g., RAS/MAPK signaling and DNA repair) that persist after treatment and could be targeted to improve patient outcomes.

### SwtichClass uncovers attenuated and escalated signalling nodes in dietary reversal phosphoproteomics

We next applied SwitchClass to a phosphoproteomic dataset profiling the effects of dietary perturbation and reversal in mice (Fig. 3A). In this experiment, animals were maintained on a normal chow diet, subjected to high-fat diet (HFD) feeding to induce insulin resistance, and subsequently switched back to chow to enable partial metabolic recovery (Fazakerley et al., 2023). These three groups represent the normal, perturbed, and reversal states, providing an experimental model of molecular adaptation and the incomplete restoration of insulin signalling. Phosphoproteome measurements were obtained from chow-fed animals under basal conditions and from all dietary groups following acute insulin stimulation (Supplementary Fig. 3A).

**Figure 3.**
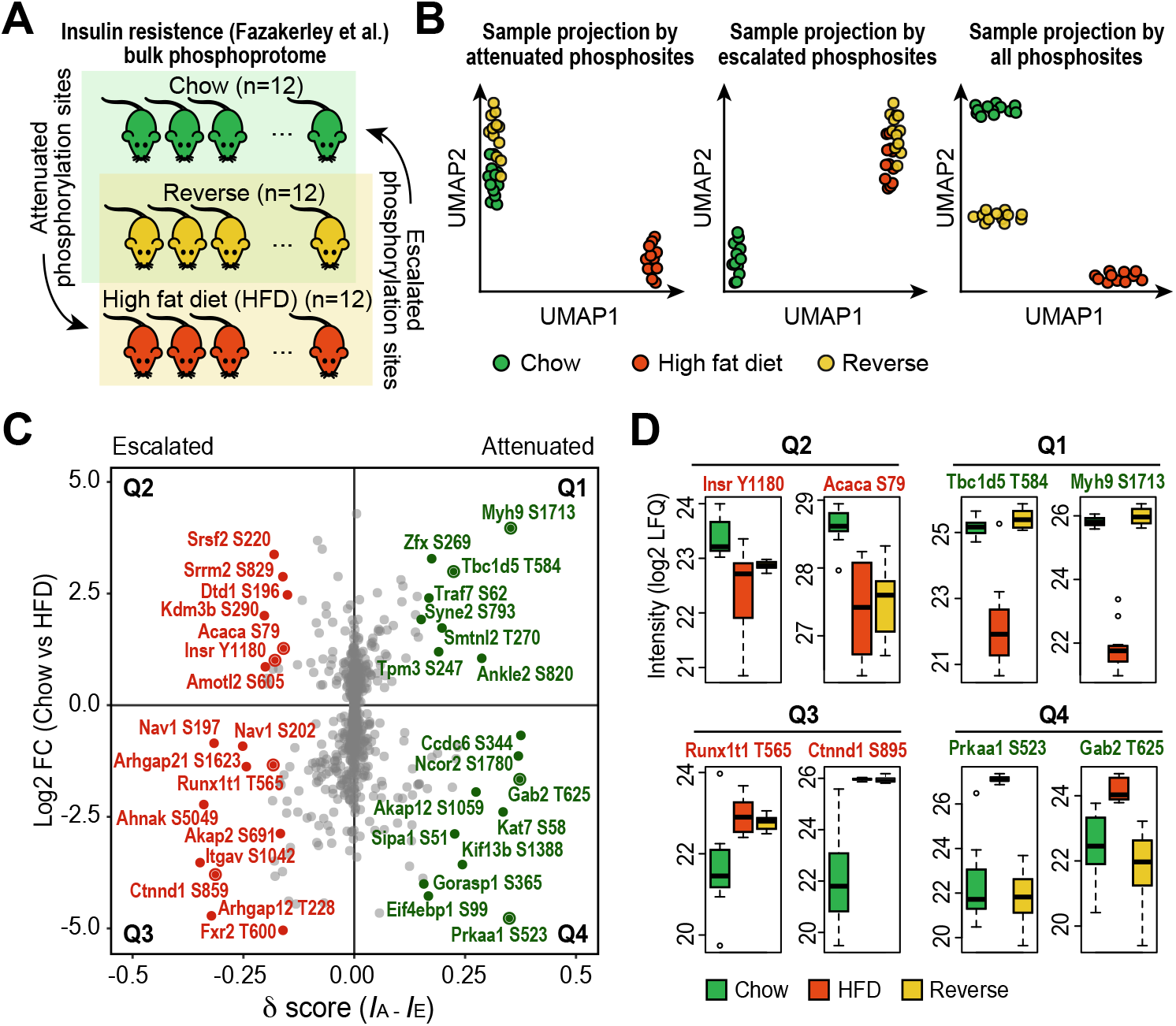
SwitchClass delineates directional phosphorylation states across dietary perturbation and reversal. (**A**) Schematic overview of dissecting phospho-proteomic profiles of mice maintained on a normal chow diet, a high-fat diet (HFD), and those subsequently switched back to the chow diet (Fazakerley et al., 2023). (**B**) UMAP visualisation of sample projections using attenuated or escalated phosphosites identified by SwitchClass, and all phosphosites. (**C**) Scatterplot of phosphosites by log2 fold change (Chow vs HFD) and differential importance scores (|*δ*|) calculated by SwitchClass. Phosphosites with high *δ* in each of the four quadrants are highlighted. (**D**) Representative sites of each of the four quadrants illustrating distinct adaptive behaviours.

Applying SwitchClass to this dataset revealed distinct directional signatures across the phosphoproteome. The method classified phosphosites according to their alignment across dietary states, identifying attenuated and escalated features that correctly grouped and separated samples in the UMAP embedding space (Fig. 3A). Across the phosphoproteome, attenuated sites were found to be related to canonical insulin-responsive nodes (e.g., AKT, mTOR and AMPK substrates) consistent with partial restoration of insulin signalling, whereas escalated sites comprised persistent insulin-receptor and metabolic phosphorylation events (e.g., INSR and ACACA substrates), highlighting incomplete reversal from HFD-induced signalling (Fig. 3C, Supplementary Fig. 3B).

Representative attenuated sites across quadrants Q1 and Q4 include Tbc1d5 T584 and Myh9 S1713 (Q1), as well as Prkaa1 S523 and Gab2 T625 (Q4) (Fig. 3D). Tbc1d5, a Rab GTPase-activating protein regulating endosomal recycling (Jia et al., 2016), showed normalisation suggesting restored vesicular transport under insulin stimulation. Similarly, Myh9 is related to GLUT4 translocations to the plasma membrane (Fulcher, Smith, Russ, & Patel, 2008) and its attenuation suggests recovery of insulin-dependent dynamics for glucose uptake. Dephosphorylation of Prkaa1 S523, an inhibitory site on the AMPK catalytic subunit (Kim et al., 2021), indicates reactivation of cellular energy sensing as insulin sensitivity improves. Finally, normalisation of Gab2, a scaffolding adaptor downstream of insulin and cytokine receptors (Nishida & Hirano, 2003), reflects reinstatement of balanced receptor-proximal signalling. Collectively, these attenuated sites reflect coordinated restoration of insulin-responsive and metabolic regulatory pathways during dietary reversal.

Conversely, persistent or escalated phosphorylation patterns were captured in quadrants Q2 and Q3 (Fig. 3D). Sites such as Insr Y1180 and Acaca S79 (Q2) remained lower in both HFD-fed and reversal animals relative to chow controls, indicating sustained suppression rather than recovery of insulin-receptor and lipogenic signalling (Cantley et al., 2019). Additional escalated responses were observed for Runx1t1 T565 and Ctnnd1 S895 (Q3). Runx1t1, a transcriptional co-regulator promoting adipogenic differentiation (Raza et al., 2022), exhibited elevated phosphorylation in the intermediate state, implying that HFD-induced adipogenic programmes persist despite dietary reversal. Ctnnd1, encoding p120-catenin, showed escalated phosphorylation at S895, consistent with ongoing cytoskeletal and junctional remodelling associated with inflammatory signalling (Kourtidis, Ngok, & Anastasiadis, 2013). These Q2-Q3 phosphosites highlight enduring repression of insulin-receptor and metabolic signalling alongside persistent stress-associated phosphorylation, underscoring that molecular recovery remains incomplete.

Together, these results show that dietary reversal produces a heterogeneous phosphoproteomic response and by separating attenuated from escalated features, SwitchClass captures this directional asymmetry in recovery and provides an interpretable framework for resolving how adaptive and residual signalling coexist during dietary intervention.

### SwitchClass delineates transcriptional states across COVID-19 disease severity

We next applied SwitchClass to a single-cell transcriptomic dataset profiling peripheral blood mononuclear cells from healthy individuals and patients with mild or severe COVID-19 (Schulte-Schrepping et al., 2020) (Fig. 4A). These three groups represent progressive immune perturbation, from homeostatic to hyperinflammatory states. Single-cell expression matrices were aggregated into pseudo-bulk profiles per cell type and patient (see Methods), and SwitchClass was used to quantify directional transcriptional alignment across severity levels.

**Figure 4.**
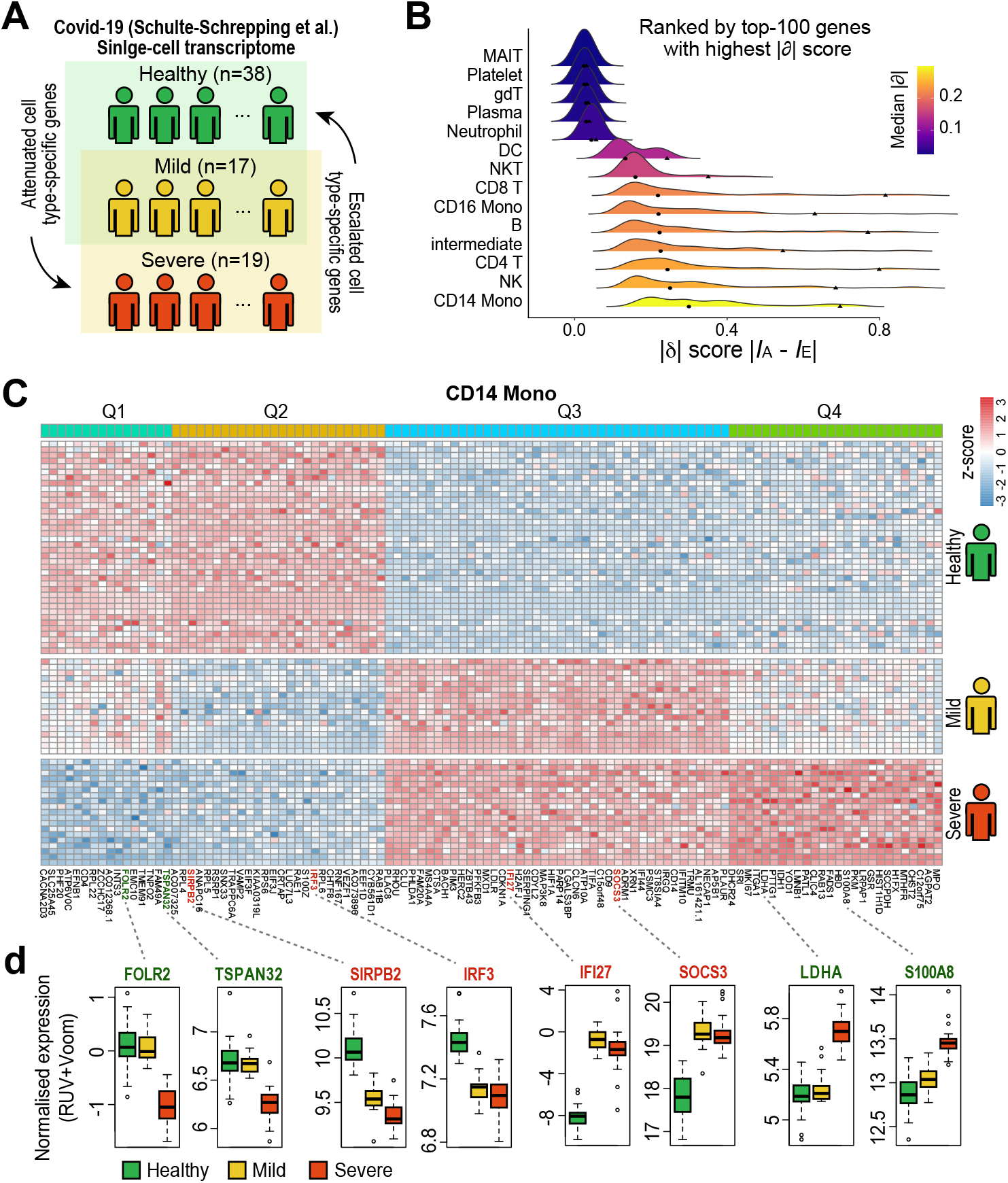
SwitchClass delineates transcriptional states across COVID-19 disease severity. (**A**) Schematic overview of dissecting single-cell transcriptome profiles from healthy individuals and patients with mild or severe COVID-19 (Schulte-Schrepping et al., 2020). (**B**) Cell-type-specific distributions of differential importance scores (|*δ*|) calculated from the top 100 genes. (**C**) Heatmap of CD14^+^ monocyte genes identified by SwitchClass across the four quadrants. (**D**) Representative attenuated (Q1, Q4) and escalated (Q2, Q3) genes in CD14^+^ monocytes in healthy individuals and mild and severe COVID-19 patients.

Across immune compartments, SwitchClass revealed heterogeneous degrees of molecular divergence (Fig. 4B). The magnitude of the differential importance score (|*δ*|) calculated from the top 100 genes varied markedly between cell types, with CD14^+^ monocytes (“CD14 Mono”) and NK cells exhibiting the largest median values, indicating strong directional transcriptional change relative to disease severity. In contrast, MAIT and platelet cells ranked lowest, reflecting minimal directional reprogramming and consistent with their restricted transcriptional activity and limited gene-expression plasticity in peripheral blood. These patterns are consistent with the previous studies (Schulte-Schrepping et al., 2020; Wilk et al., 2020) and highlight that directional molecular divergence primarily arises within the highly responsive innate compartments, particularly monocytes and NK cells.

In CD14^+^ monocytes, which displayed the most pronounced directional signatures, SwitchClass stratified genes into four quadrants based on their *δ* values and gene expression patterns (Fig. 4C). Representative attenuated transcripts included FOLR2 and TSPAN32 (Q1) as well as LDHA and S100A8 (Q4) (Fig. 4D). FOLR2, encoding a folate receptor enriched by macrophages (Puig-Kröger et al., 2009), and TSPAN32, a tetraspanin implicated in negative regulation of immune activation and cytokine signalling (Fagone, Mangano, & Nicoletti, 2024), were both reduced in severe disease but maintained in healthy and mild cases, consistent with suppression of anti-inflammatory and regulatory monocyte programs under hyperinflammatory conditions. Similarly, lower expression of LDHA, a key glycolytic enzyme driving inflammatory metabolic reprogramming (Arra et al., 2020), and S100A8, a heterodimer constitutively expressed in monocytes and associated with inflammation (S. Wang et al., 2018), in healthy and mild patients reflected attenuation of hyperinflammatory and glycolytic pathways. Conversely, transcripts in quadrants Q2 and Q3 exhibited divergent patterns between healthy individuals and those from mild and severe patients (Fig. 4D). SIRPB2, a gene that modulates cytokine production and innate immunity (Visser et al., 2025), and IRF3, a transcription factor crucial for the innate immune response to viral infections (Grandvaux et al., 2002), were reduced in both mild and severe disease relative to healthy controls, suggesting blunted antiviral responsiveness in hyperinflammatory monocytes. In contrast, IFI27, an interferon-inducible gene associated with antiviral responses (Shojaei & McLean, 2025), and SOCS3, a suppressor of cytokine signalling elevated during chronic inflammation (Carow & Rottenberg, 2014), exemplify alleviated immune responses.

Similarly, in NK cells, SwitchClass identified directional transcriptional patterns that dissect the attenuated and alleviated immune responses and interferon activities (Supplementary Fig. 4A,B). Example Q1 genes, such as NFATC3, an inflammation-responsive TF (Urso et al., 2011), and MYO1F, exclusively expressed in immune cells (Cui et al., 2025), and Q3 genes, such as GSTP1, which regulate oxidative stress (Mian et al., 2016), and JAK3, essential for the growth and maturation of NK cells (Cornejo, Boggon, & Mercher, 2009), Q4 were attenuated in healthy and mild cases compared to severe cases. In contrast, Q2 genes GIMAP1, essential for NK cell survival and function (Limoges, Cloutier, Nandi, Ilangumaran, & Ramanathan, 2021), and SIGLEC7, an immune checkpoint receptor (Zheng et al., 2020), and Q3 genes MX1, a key gene for innate antiviral response (Ciancanelli, Abel, Zhang, & Casanova, 2016), and ISG15, a gene that codes a ubiquitin-like protein for inhibiting viral replication (Kang, Kim, & Jeon, 2022), were more similar between mild and severe cases compared to healthy. Together, these features reveal a decoupling of homeostatic regulatory programs and inflammatory activity in CD14^+^ monocytes and NK cells characteristic of COVID-19 severity.

## Discussion

Biological systems rarely transition cleanly between conditions such as health and disease. Instead, they often pass through intermediate states that show partial recovery or persistent molecular reprogramming. In this study, we introduced SwitchClass, a label-switch classification framework that quantifies directional molecular behaviour in intermediate states and explicitly distinguishes features that attenuate towards the reference state from those that escalate towards the perturbed state. Across controlled simulations and case studies spanning various biotechnologies (proteome, phosphoproteome, and transcriptome modalities) and biological systems, SwitchClass robustly separated attenuated from escalated features for recovering directionally informative molecular profiles. These benchmarks establish that in the presence of intermediate states, explicitly modelling directional alignment provides statistical and interpretative advantages over conventional two-group contrast analyses.

Methodologically, SwitchClass differs from prior strategies in two aspects. First, the label-switch design converts the intermediate state problem into two complementary binary classifications that share the same features but invert its grouping. The contrast of feature-importance vectors from these classifiers yields a single directional score, *δ*, that is straightforward to interpret and to threshold. While random forest was employed in the current model, the conceptual framework is agnostic to the choice of classifier and importance metric, allowing practitioners to employ linear models for interpretability or other non-linear models for non-linear effects, and to estimate stability via bootstrapping.

From a biological perspective, the quadrant stratification (Q1–Q4) situates each feature within both direction (attenuated vs escalated) and regulation (activation vs repression), enabling downstream interpretation such as pathway enrichment and network analyses. For example, escalated and activated phosphosites can nominate kinase activities that remain pathologically high after dietary reversal, whereas attenuated and repressed transcripts in COVID-19 may indicate restoration of antiviral defences. Such directed summaries are useful for prioritising targets for intervention or for monitoring treatment response with minimal panels.

There are several limitations that future extension may address. First, the current implementation assumes three biological states. Many studies include richer time courses or multiple intermediate arms; while SwitchClass could be extended by fitting multiple label-switch contrasts (e.g., against early/late intermediates) and aggregating *δ* scores, principled multi-state generalisations are an avenue for future work. Secondly, feature importance from non-linear models can be biased by correlations; we partially mitigate this by bootstrap aggregation, yet complementary approaches (e.g., conditional importance or knockoff-based attribution) may improve identifiability. Thirdly, class imbalance and batch structure can affect classifier training. In our applications we used donor-level aggregation and standard normalisation to minimise confounding, but explicit incorporation of mixed-effects or domain-adaptation may be effective alternative approaches for complex data structure.

Finally, the framework complements, rather than replaces, existing differential and pathway tools. We envisage a typical workflow in which SwitchClass first assigns directional labels to features, followed by enrichment, network inference, and causal hypothesis testing targeted to the relevant quadrants. This modularity should make the method easy to adopt across omic modalities and experimental settings.

## Conclusion

SwitchClass presents a general strategy for dissecting directional molecular behaviour in systems that exhibit partial alteration. By reframing classification-based feature importance through a label-switch design, SwitchClass quantifies attenuated and escalated molecular profiles using continuous and interpretable scores. The framework enable analyses across transcriptomic, proteomic, and phosphoproteomic, revealing mechanisms of adaptation and maladaptation and providing a foundation for comparative and translational studies.

## Methods

### SwitchClass computational framework

SwitchClass was developed to quantify directional molecular behaviour across systems comprising three biological states that represent normal reference, perturbed or disease conditions, and intermediate or reversal conditions. The method is based on a label-switch strategy that uses complementary binary classifications using models that are capable of ranking feature importance to capture the differential contribution of molecular features to each direction of change. This design enables systematic detection of features whose intermediate profiles either match to the reference state (attenuated) or are aligned with the perturbed state (escalated).

Specifically, in the current implementation, two random forest classifiers (Breiman, 2001) are trained under inverted label configurations. In the first, termed the reference-aligned configuration, samples from the normal reference and intermediate groups are pooled and classified against the perturbed group. In the second, termed the perturbation-aligned configuration, the intermediate and perturbed samples are grouped and classified against the normal reference. Each model yields a feature-importance vector, *I*_A_ from the reference-aligned model and *I*_E_ from the perturbation-aligned model, quantifying the contribution of each feature to class discrimination. The directional importance score for feature *i* is defined as

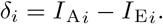

Positive *δ*_*i*_ values indicate that a feature contributes more strongly under the reference-aligned model and therefore exhibits attenuated behaviour, whereas negative *δ*_*i*_ values indicate stronger contribution under the perturbation-aligned model and therefore escalated behaviour. The absolute magnitude |*δ*_*i*_| reflects the strength of directional alignment.

To facilitate interpretation, features are stratified into four quadrants (Q1-Q4) according to the sign of *δ*_*i*_ and the direction of fold-change between normal references and perturbed conditions. This classification delineates attenuated versus escalated features while distinguishing activation and repression patterns within each category. For stability estimation, the training procedure is repeated across 20 bootstrap resamples of samples, and the median *δ*_*i*_ across replicates is reported as the final estimate. The resulting directional importance scores are used to identify molecular subsets underlying the directional organisation of intermediate states.

### Simulation benchmark

To evaluate the performance and robustness of SwitchClass, we generated simulated datasets in which the underlying directional behaviour of each feature was predefined. Each dataset comprised three groups representing normal reference, intermediate, and perturbed ‘disease’ states, corresponding conceptually to the configuration of the SwitchClass framework. A total of 1,000 features were simulated from multivariate normal distributions with controlled variance *X* ∼ *N* (*µ*_*s*_, 1), partitioned into four behavioural categories. Specifically, 50 features were defined as *attenuated* with mean pattern (0, *α*, 0) of the three states, 50 as *escalated* with pattern (0, *α, α*), 200 as *intermediate-specific* with pattern (0, 0, *α*), and 700 as *null* features with no systematic differences (0, 0, 0).

We systematically varied the mean separation between the normal reference and perturbed groups (effect size, *α*) and the number of samples per group (*n*) to examine the influence of signal strength and sample size on method performance. Effect sizes were drawn from *α* ∈ {0.6, 1.0, 1.6}, corresponding to ∼ 1.5-, ∼ 2-, and ∼ 3-fold differences, and sample sizes from *n* ∈ {10, 20}, covering a range of realistic experimental settings. For each simulation, we computed directional importance scores (*δ*) using SwitchClass and compared its ability to recover the known feature classes with a conventional pairwise differential expression approach that compares intermediate vs perturbed and normal vs intermediate groups for detecting attenuated and escalated features, respectively, using the limma R package (Ritchie et al., 2015).

Performance was evaluated using the area under the precision-recall curve (AUPRC) calculated with the PRROC package (Grau, Grosse, & Keilwagen, 2015). For each *α* and *n* combination, five independent repetitions (*B* = 5) were generated, and mean ± 95% confidence intervals were computed from repetitions. Visualisation of sample-level structure was achieved using UMAP embeddings derived from attenuated, escalated, intermediate-specific, or null feature subsets, or full features, demonstrating how directional separation of samples was recovered under different simulation conditions.

### Experimental datasets preprocessing and differential analysis

We demonstrate the utility of SwitchClass on three experimentally derived datasets capturing reversible or progressive molecular adaptations in distinct biological contexts. The first dataset is a human plasma proteomic dataset derived from a longitudinal colorectal cancer cohort (Y. Li et al., 2024). For this dataset, raw protein level intensity values were obtained from the original study and subset to include healthy controls, oxaliplatin-sensitive pre-treatment samples, and post-treatment samples from the stable disease subgroup (SSG). Protein intensities were log2-transformed and filtered to retain those quantified in at least 10% of samples within each group. Missing values were imputed using the PhosR package (Kim et al., 2021). Differential expression analysis was performed using limma, comparing healthy controls to pre-treatment samples. Proteins with an adjusted *p*-value *<* 0.05 and an absolute fold change of 1.5 were retained as differentially expressed proteins. This curated dataset captures a directional proteomic response across health, disease, and therapeutic stabilization, and was used for all downstream SwitchClass analyses.

The second dataset is generated from a mouse adipose phosphoproteomic experiment profiling dietary perturbation and reversal (Fazakerley et al., 2023). Previously processed phosphosite intensities (log2 LFQ) by PhosR (Kim et al., 2021) were obtained from the original study. These values were derived from adipose tissues collected under chow basal conditions, and from chow, HFD, and reversal groups following acute insulin stimulation. To focus on insulin-regulated signalling, we first performed differential analysis within chow animals (insulin vs basal) and retained phosphosites that were significantly insulin-responsive (Benjamini–Hochberg FDR *<* 0.05) with an absolute fold change of 1.5. The resulting insulin-responsive set was used for all downstream analyses on insulin-stimulated samples. This dataset provides a controlled model for examining phosphorylation events that normalise or remain altered during the reversal of diet-induced insulin resistance.

The third dataset is generated from a single-cell transcriptomic study of peripheral blood mononuclear cells (PBMCs) across COVID-19 disease severity (Schulte-Schrepping et al., 2020). Count matrices from the original study were obtained. To ensure sufficient sampling depth per donor and cell type, we first retained only patient– cell type pairs containing at least 20 cells. Pseudo-bulk counts were then constructed by summing gene counts for each patient–cell type combination. We next restricted to cell types with adequate patient representation in each disease group by requiring, for that cell type, at least five patients per group across healthy, mild, and severe. Within each retained cell type, patient-level counts were TMM-normalised and transformed with Voom (Law, Chen, Shi, & Smyth, 2014). To mitigate unwanted variation while protecting the severity effect, we applied RUV normalisation (Gagnon-Bartsch & Speed, 2012) on the voom-transformed matrix. Differential expression was then performed between healthy and severe patients using limma to select for the top 1000 genes that vary in expression between the two conditions. These steps enable robust modelling of cell-type-specific transcriptional variation at the donor level. Together, these datasets serve as examples of molecular adaptation and incomplete recovery at signalling and transcriptional levels.

## Data availability

No new experimental data were generated in this study. The colorectal cancer datasets were obtained from (Y. Li et al., 2024), the insulin resistance phosphoproteomic dataset from (Fazakerley et al., 2023), and the COVID-19 single-cell transcriptomic dataset from (Schulte-Schrepping et al., 2020).

## Author contributions

P.Y. conceived and supervised the study. P.Y. and D.X. performed the analysis and drafted the paper. A large language model (ChatGPT-5, OpenAI) was used to assist with copy-editing during manuscript preparation. All authors read, edited, and approved this work.

## Acknowledgement

We thank the colleagues from Computational Systems Biology Unit for their support and intellectual engagement. This work was supported by a Robinson Fellowship (G200734) to P.Y.

## Supplementary Figures

**Supplementary Figure 1.**
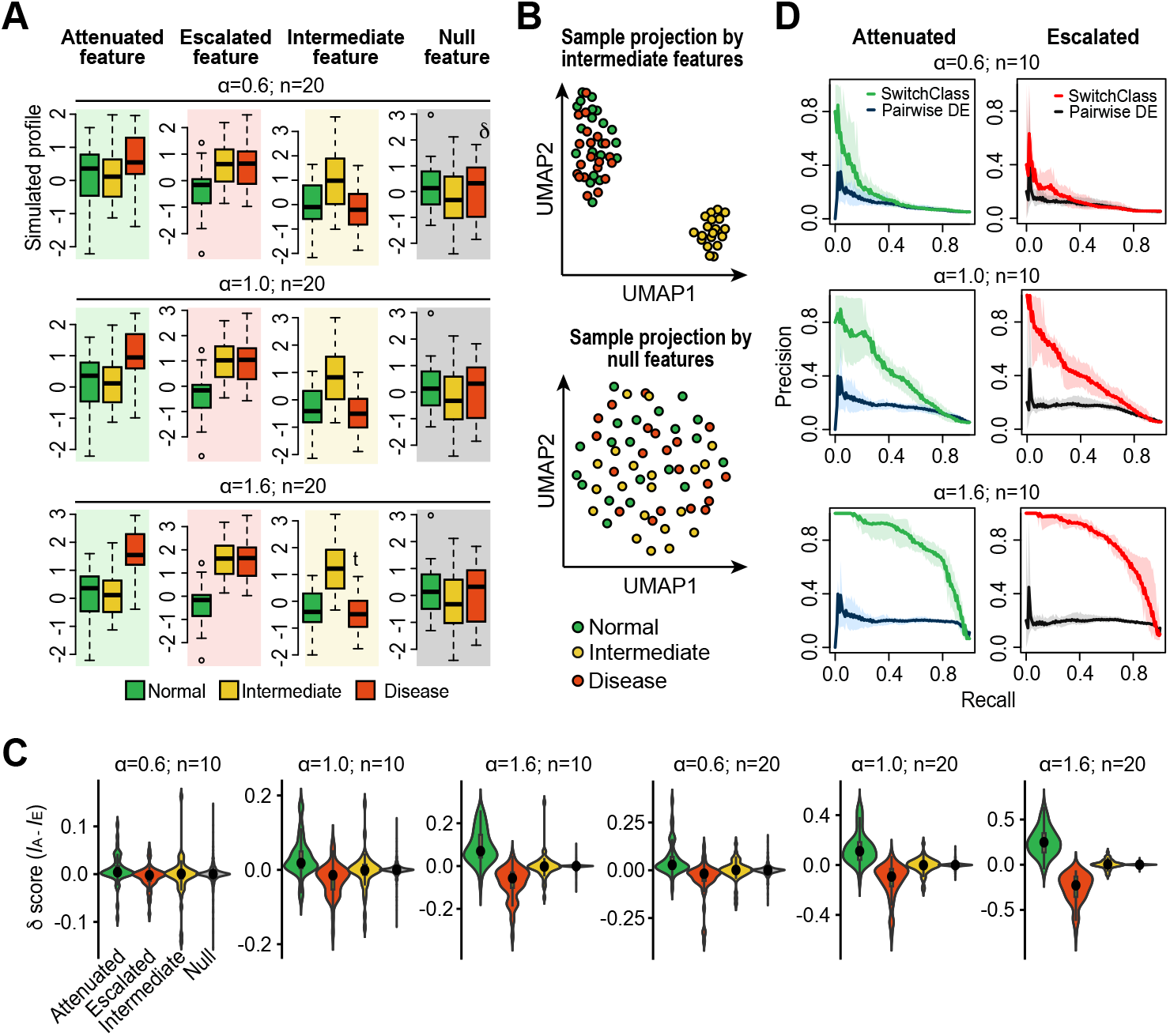
Simulations and benchmarks of SwitchClass under varying effect sizes and sample sizes. (**A**) Examples of simulated expression profiles of attenuated, escalated, intermediate, and null features across normal, intermediate, and disease states for three effect sizes (*α* = 0.6, 1.0, 1.6) and sample size of *n* = 20. (**B**) UMAP projections of samples using intermediate and null features. (**C**) Distributions of differential importance scores *δ* = *I*_A_ − *I*_E_ from SwitchClass across feature classes, effect sizes (*α* = 0.6, 1.0, 1.6), and sample sizes (*n* = 10, 20). (**D**) Precision-recall performance of SwitchClass in detecting attenuated and escalated features compared with pairwise differential expression (DE) across varying effect sizes (*α* = 0.6, 1.0, 1.6) and a sample size of *n* = 10 for each state. Shaded areas denote 95% confidence intervals derived from bootstrap resampling across simulation replicates.

**Supplementary Figure 2.**
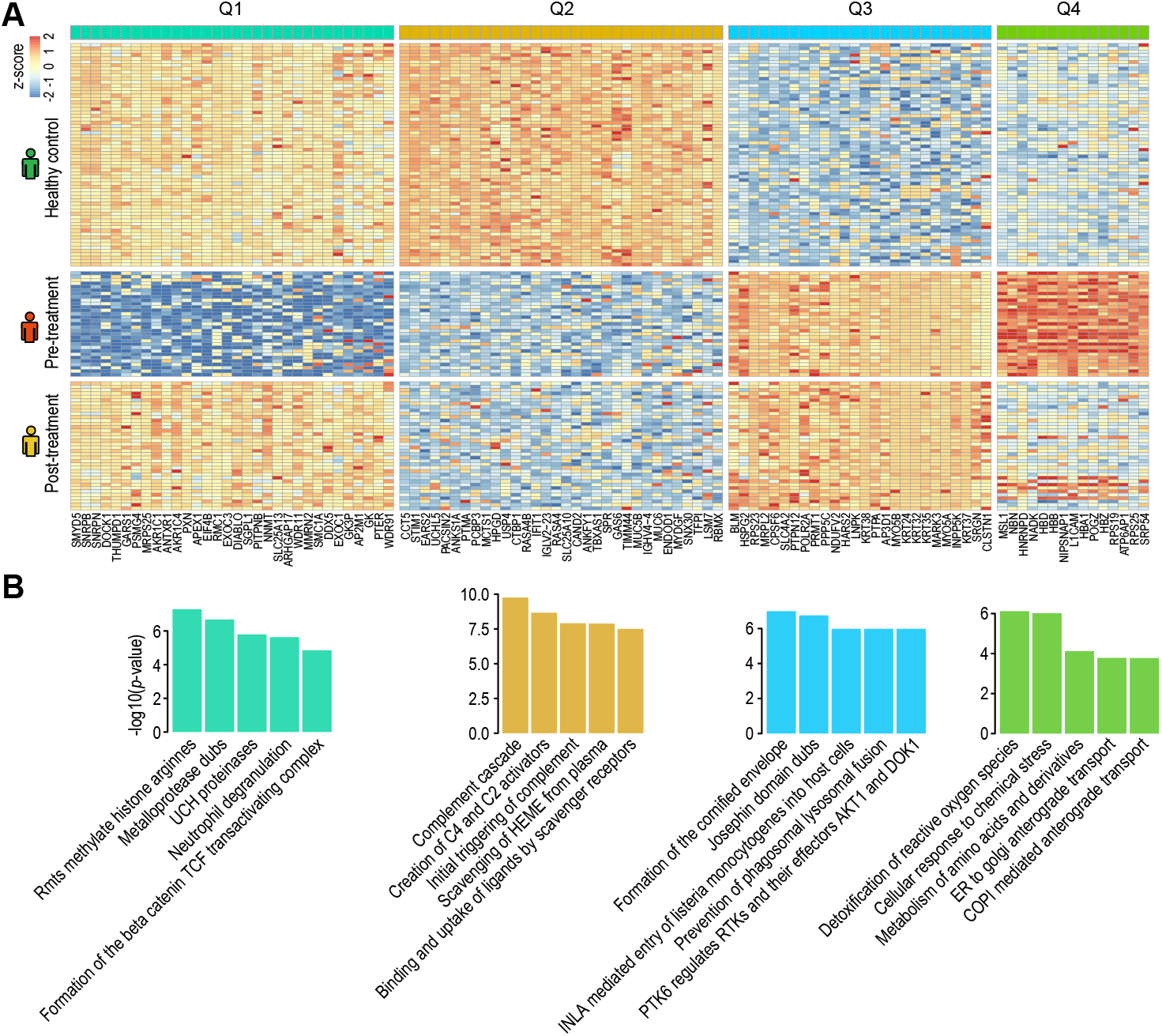
Plasma proteomic expression and SwitchClass identified attenuated or escalated proteins in colorectal cancer and after its treatment. (**A**) Heatmap of proteins identified by SwitchClass as attenuated (Q1, Q4) or escalated (Q2, Q3) across healthy controls, and in colorectal cancer pre-treatment and post-treatment samples. (**B**) Pathway enrichment analysis of quadrant-specific proteins using the Reactome database.

**Supplementary Figure 3.**
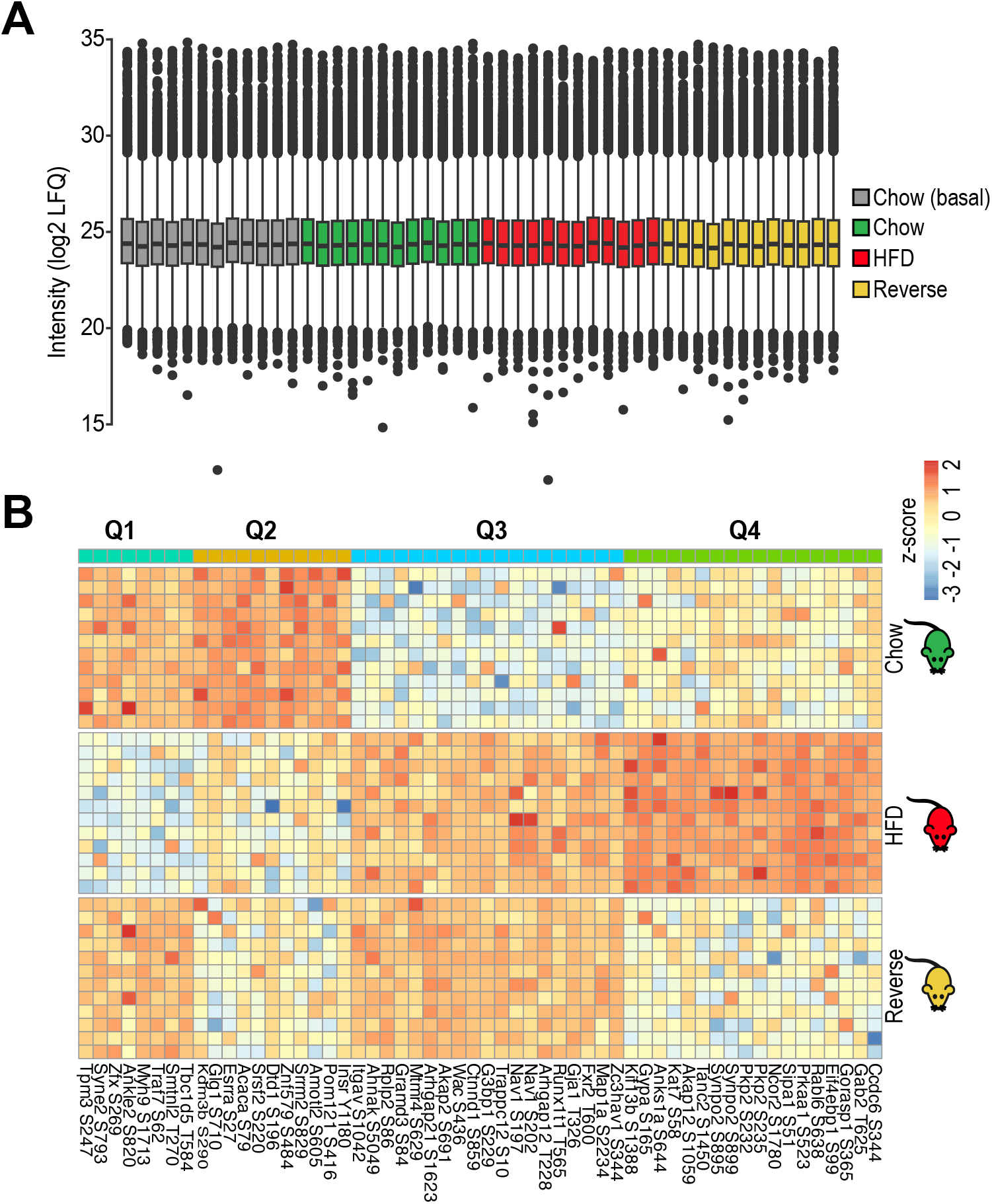
Phosphoproteomic data distribution and Switch-Class identified attenuated or escalated phosphosites across dietary perturbation and reversal. (**A**) Boxplots showing the distribution of log_2_ LFQ intensities across all samples, including basal (chow) and insulin-stimulated chow, high-fat diet (HFD), and reverse groups. (**B**) Heatmap of phosphosites identified by SwitchClass as attenuated (Q1, Q4) and escalated (Q2, Q3) in mice from Chow, HFD, and reverse states.

**Supplementary Figure 4.**
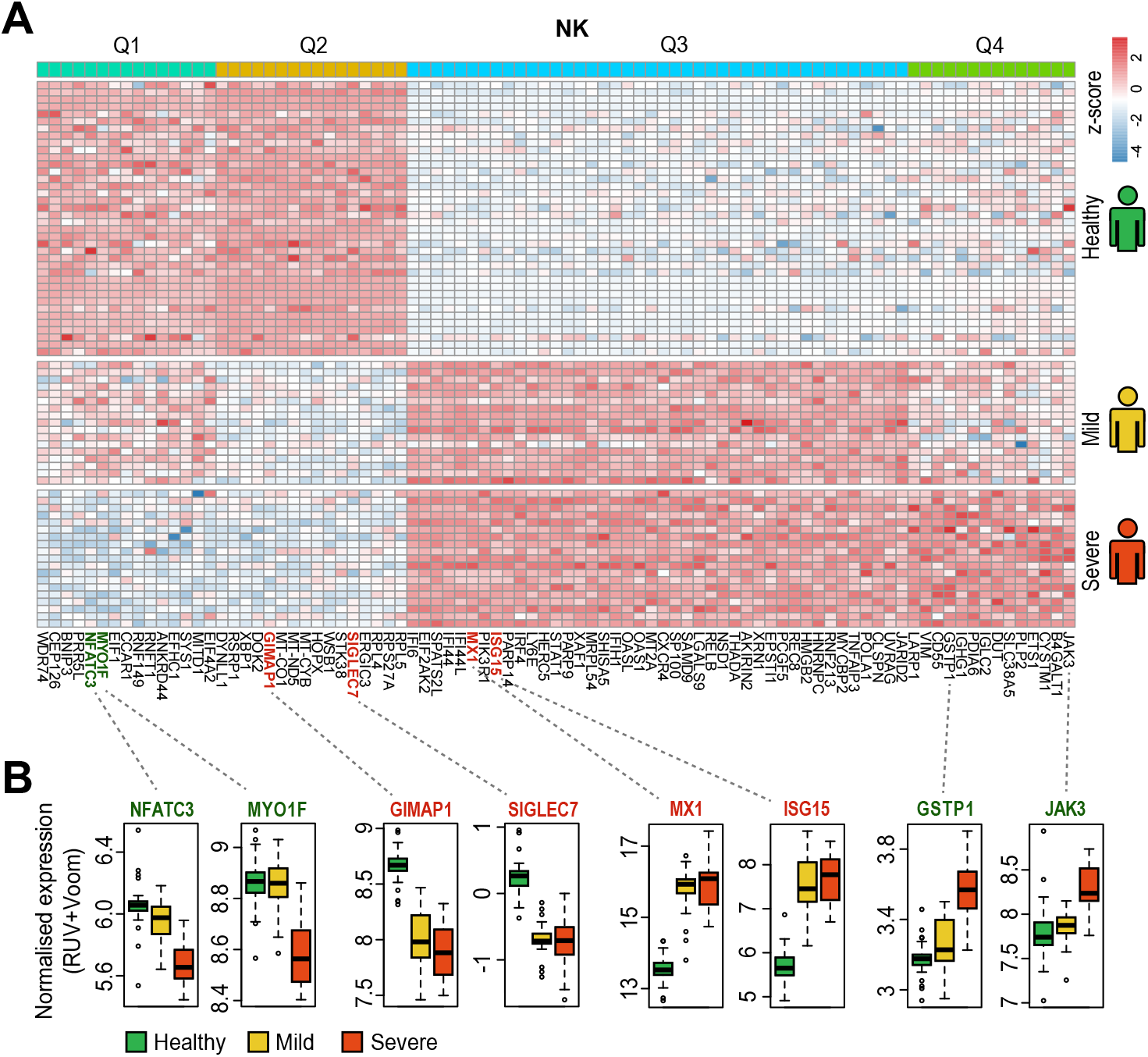
SwitchClass identified attenuated or escalated genes in NK cells in healthy individuals, and mild and severe COVID-19 patients. (**A**) Heatmap of expression profiles of genes identified by SwitchClass across the four quadrants in NK cells. (**B**) Representative attenuated (Q1, Q4) and escalated (Q2, Q3) genes in NK cells in healthy individuals and mild and severe COVID-19 patients.

